# Adolescent-onset and adult-onset schizophrenia: reduced ribosomal protein expression via mTOR signalling in patient-derived olfactory cells

**DOI:** 10.1101/2020.08.26.267930

**Authors:** Yichen Li, Melanie Föcking, Alexandre S. Cristino, Jane English, Gerard Cagney, Anthony James, David Cotter, Francis G. Szele, Alan Mackay-Sim

**Author notes:** These authors contributed equally (Yichen Li, Melanie Föcking, Alexandre S. Cristino). Corresponding authors. AMS phone: ±61 407 756 740;, FGZ.

## Abstract

Schizophrenia is a heterogeneous disorder associated with many genetic and environmental risk factors that could affect brain development. It is unknown whether adolescent-onset and adult-onset schizophrenia have similar aetiology. To address this we used discovery-based proteomics to find proteins differentially expressed in olfactory neurosphere-derived cells from adolescents with schizophrenia compared to age- and gender-matched healthy controls. Of 1638 proteins identified, 241 were differentially expressed in patient cells, with significant down-regulation of ribosomal and cytoskeletal proteins, and dysregulation of protein synthesis pathways. We then re-analysed our previous adult-onset proteomic data to compare directly with adolescent-onset protein expression. Schizophrenia-associated protein expression in adult-onset patients was remarkably similar to adolescent-onset patients. To increase sample size and power we combined the two datasets for a bioinformatic meta-analysis. Schizophrenia-associated protein expression indicated significant downregulation of the mTOR signalling pathway, which regulates protein synthesis, indicated by the reduced expression of all ribosomal proteins and other mTOR-dependent proteins: RPS6, VIM, LDHB and PPP2R1A. A protein-protein interaction network built from differentially expressed proteins in the combined dataset was significantly associated with schizophrenia-associated risk genes and with proteins regulating neural stem cell differentiation, cell adhesion and growth cones in the developing brain. This study demonstrates that despite the divergent age of onset, the proteomes of olfactory neural stem cells of adolescent- and adult-onset patients are remarkably similar. The dysregulated proteins in patient cells form a tightly interconnected protein-protein interaction network associated with mTOR signalling, protein translation, neurogenesis and axon growth - all key components of brain development.

## Introduction

Adolescent-onset schizophrenia is severe, with high morbidity and mortality. Treatment resistance is common, and prognosis often poor ^1^. Why some patients with schizophrenia present with disease during adolescence and others during adulthood is poorly understood. Adolescent-onset, and adult-onset schizophrenia share genetic and environmental risk factors which ultimately alter protein expression and disrupt neurodevelopment ^2, 3^. The challenge for the field is to link the molecular and cellular processes that cause schizophrenia with neurodevelopment and the complex underlying genetic risk ^4^. Patient-derived stem cells are a means to identify how schizophrenia-associated differences in molecular expression contribute to changes in cell development or function to cause schizophrenia.

Patient-derived stem cells from the olfactory mucosa are an accessible source of primary cultured cells that can be stored frozen and expanded readily. These olfactory neurosphere-derived cells, “ONS cells”, are a convenient model of early stem cell-based developmental events ^5, 6^. ONS cells show very robust differences between patients with adult-onset schizophrenia and matched healthy controls in gene transcription, DNA methylation and multiple cell functions ^5, 7–10^. Adult onset patient-derived ONS cells have significantly reduced expression of ribosomal proteins and dysregulated expression of several proteins involved in protein synthesis ^11^. A protein translation assay demonstrated that patient-derived cells, compared to control-derived cells, had a significantly reduced rate of protein synthesis^11^.

The aim of the present study was to reveal the proteome of adolescent-onset patient-derived ONS cells for comparison with adult-onset patient-derived ONS cells, in part to understand whether the aetiology of adolescent-onset schizophrenia is similar to, or different from, adult-onset schizophrenia. This study is multi-faceted: 1) a discovery-based, proteomic analysis of ONS cells from adolescent-onset schizophrenia patients and age-matched healthy controls; 2) a re-analysis of the raw data from our proteomic analysis of adult-onset patients and controls^11^; 3) comparisons of the proteomes of adolescent and adult patients and controls to test the overlap of differentially expressed proteins; 4) a bioinformatic meta-analysis of the combined data from adolescent- and adult-onset studies; 5) construction of a protein-protein interaction network based on the proteins differentially expressed in the combined dataset; and 6) re-analysis of published proteomics datasets concerned with neurogenesis and axon growth in the developing mouse brain to determine their overlap with proteins in the protein-protein interaction network in schizophrenia patient cells.

## Materials and Methods

### Participants

Adolescent patients were diagnosed according to DSM-V criteria. To assess psychosis, intelligence and cognitive functioning, participants were screened with the structured psychiatric interview (K-SADS-PL) ^12^ and the psychometric interview ^13^ at the Highfield Unit, Warneford Hospital, Oxford. Control subjects who showed emotional, behavioural or psychiatric disorders were excluded. Participants with any of the following conditions were also excluded: infectious or other nasal conditions, coagulopathies, known systemic medical diseases and infections, neurological disorders including significant head injury, learning disability, substance abuse or poor understanding of English. All eligible participants, or in the case of younger participants (<18 years old) their parents, gave signed consent for participation in the study. The study was approved by the South Central-Oxford A Research Ethics Committee (IRAS no 104383). Patient and control demographic data are shown in Table 1. Adult patients and age- and sex-matched controls were those recruited as described previously ^5, 11^.

**Table 1.**
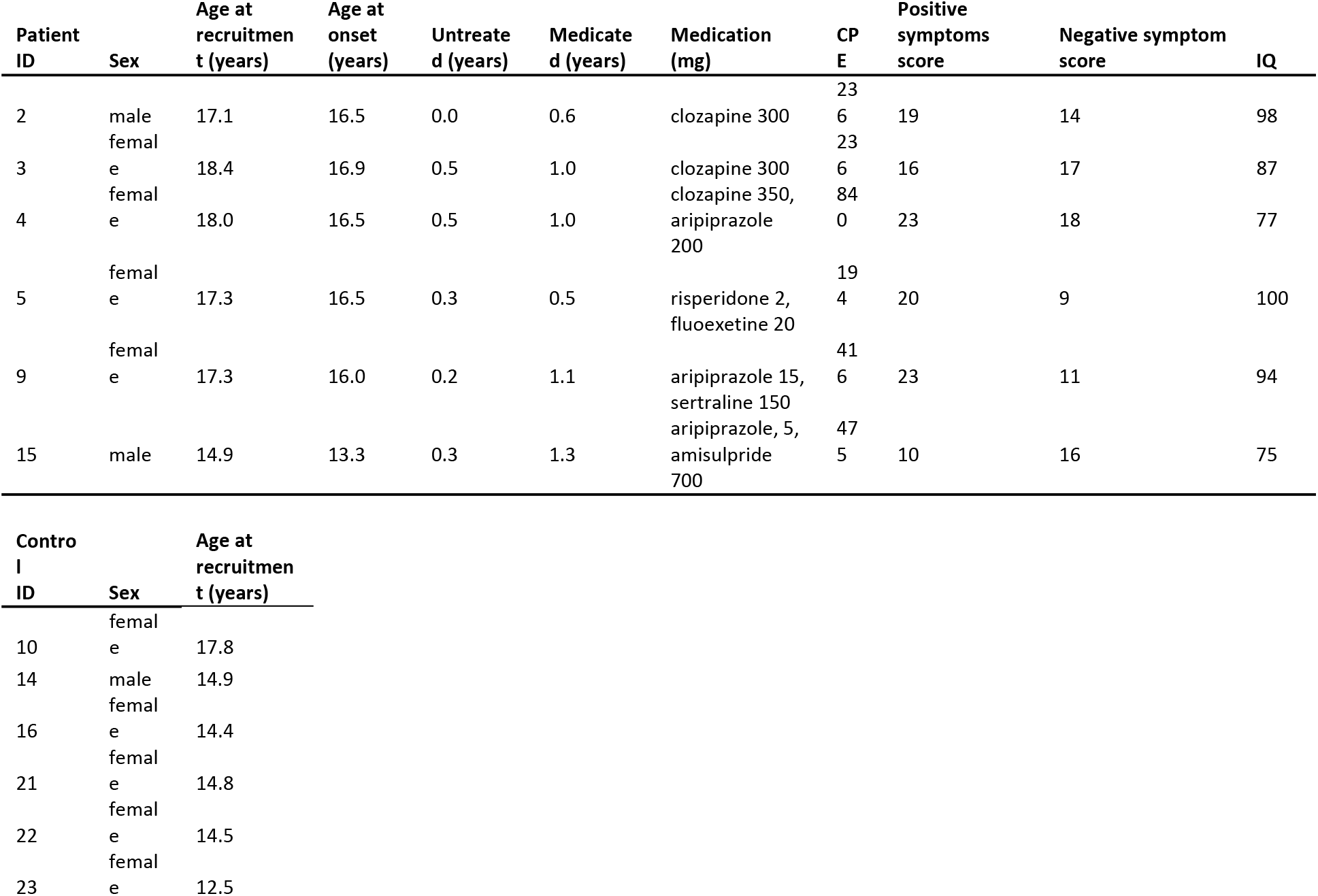
Biometrics of adolescent-onset patients and healthy controls.

### Cell culture

Small biopsies were removed from the olfactory mucosa on the dorsal posterior nasal septum under local anaesthesia, placed in primary culture medium (Dulbecco’s Modified Eagle Medium, DMEM/HAM F12 with GlutaMax, Gibco 31331; 10% fetal bovine serum, FBS, Gibco 10270-106; and 1% Penicillin-Streptomycin, P/S, Life Technologies 15140-122) and grown under standard conditions (37°C, 5% CO_2_,) as previously described ^14, 15^. Olfactory neurospheres were generated from primary cultures according to published procedures ^5, 15^. Briefly, primary olfactory cells were passaged into neurosphere formation medium (DMEM/HAM F12 with GlutaMax; 1% P/S; 1% Insulin-Transferrin-Selenium, Gibco 41400; 75 ng/ml epidermal growth factor, EGF, Chemicon GF144; and 75 ng/ml fibroblast growth factor 2, Chemicon GF003), which was refreshed daily. After a few days this produces free-floating spheres of cells (neurospheres) consisting of multipotent stem cells ^5, 6^. Olfactory neurosphere-derived cells (ONS cells) were generated by placing the neurospheres without dissociation in primary culture medium. After 4 days, ONS cells migrated out of the spheres and reached 90% confluency at which point they were passaged and maintained in primary culture medium, as described ^5^.

### Mass spectrometry

Proteomic analysis by label free LC-MS/MS analysis was carried out as previously described ^11, 16^. Protein samples were prepared from each patient and control ONS cell culture by washing them in PBS, treating with lysis buffer (proteinase inhibitor, Roche complete, 0469315900, in 10 ml of 1 M triethylammonium bicarbonate (TEAB, Sigma T7408), and immediately removing the cells with a cell scraper. Total lysates were centrifuged at 24,000 x g and 4°C for 30 min and the proteins were stored at −80°C. Protein concentration was determined using a Bradford Assay ^17^ according to the manufacturer’s (BioRad) instructions. Protein from each ONS cell line (n=12) was run on a Thermo Scientific Q Exactive mass spectrometer connected to a Dionex Ultimate 3000 (RSLCnano) chromatography system. Tryptic peptides (5μl of digest) from each sample were loaded onto a fused silica emitter (75 μm ID, pulled using a laser puller (Sutter Instruments P2000), packed with UChrom C18 (1.8 μm) reverse phase media (nanoLCMS Solutions LCC) and was separated by an increasing acetonitrile gradient over 90 minutes at a flow rate of 250 nL/min. The QC standard was injected 3 times at the beginning of the MS study to condition the column, and after every ten injections throughout the experiment to monitor the MS performance. The mass spectrometer was operated in data dependent TopN 8 mode, with the following settings: mass range 300-1600Th; resolution for MS1 scan 70000; AGC target 3e6; resolution for MS2 scan 17500; AGC target 2e4; charge exclusion unassigned, 1; dynamic exclusion 40 s. Label-free quantification (LFQ) was performed with Max Quant (V1.5.5.1; www.maxquant.org) as described ^18–20^. Protein and peptide FDR’s were set to 0.01, and only proteins with at least two peptides (one uniquely assignable to the protein) were considered as reliably identified.

### Bioinformatics and data analysis

Detected proteins were assigned to UniProt IDs and official gene names obtained from the Human Genome Nomenclature Committee database (www.genenames.org). Label-free quantification (LFQ) intensity values were log2 transformed to reduce skewness and assure the data is normally distributed. Data imputation was used to replace missing values by the average value estimated from normal distribution. The log2 transformed values were normalized by subtracting the median LFQ intensity value for all proteins per sample. Student’s t-test was used to identify proteins differentially expressed between schizophrenia patients and controls. Proteins were considered differentially expressed if the patient-control difference in expression was significantly different (p<0.05, not corrected for multiple testing). To compare the proteomes of adolescent- and adult-onset schizophrenia patients, we normalized the published proteome data from adult onset schizophrenia cohort ^11^ using the same procedure, to identify the 900 proteins detected in both proteomes (adolescent: 6 patients and 6 controls; adult: 9 patients and 9 controls). When combining adolescent and adult data sets, batch effects were minimized by subtracting the median of the normalized LFQ intensity values for each protein across all 30 samples (15 schizophrenia patients and 15 controls). Ingenuity pathway analysis (IPA; www.ingenuity.com) was performed on statistically significant proteins (p<0.05) using Canonical Pathway analysis. The enrichment of affected pathways was calculated using Fisher’s Exact Test and multiple hypothesis correction was based on the Benjamini-Hochberg correction method.

The ONS-SZ protein-protein interaction (PPI) network was built from ‘seed’ proteins and their first-degree interacting neighbours based on experimental data (at least two published studies) as described in the BioGRID database (thebiogrid.org) following the same methods as previously published ^21^. To test the association between PPI networks we used the standardized Z-score estimated by a binomial distribution described elsewhere ^21^. Distributions of network properties (degree, average path length, clustering coefficient) were generated using simulation. Fifty proteins from the 164 differentially expressed proteins were randomly selected and used as “seeds” to build a network from which the three network properties were calculated for each protein in the set. This was repeated 1000 times. A “control” simulation iteratively drew fifty proteins at random from those in the neurodevelopmental disease AXAS-PPI network previously published ^21^.

## Results

### Participants

The mean ages of the adolescent patients and controls were 15.0±0.5 (n=6) and 14.8±0.7 (n=6), respectively. The mean age of disease onset in the patients was 16.0±0.5, with 0.3±0.1 untreated years and 1.2±0.2 years of medication. The average “chlorpromazine equivalents, CPE”, a metric used to compare medications 22, 23, was 399.5±99.3 among the patients. There were 2 males and 4 females in the patient group and 1 male and 5 females in the controls (Table 1); none smoked. The mean ages of the adult patients and controls were 38.0±4.0 (n=9) and 36.0±4.3 (n=9), respectively and average CPE was 522.1±107.1^11^. All adult participants were males; 5 patients and 2 controls smoked^11^.

### Protein expression distinguished adolescent-onset patient from control ONS cells

We measured the relative expression levels of 1638 proteins identified in the 12 ONS cell samples (Supplementary Table 1). A principal component analysis of the protein expression levels separated the patient and control samples into two distinct clusters (Figure 1A).

**Figure 1.**
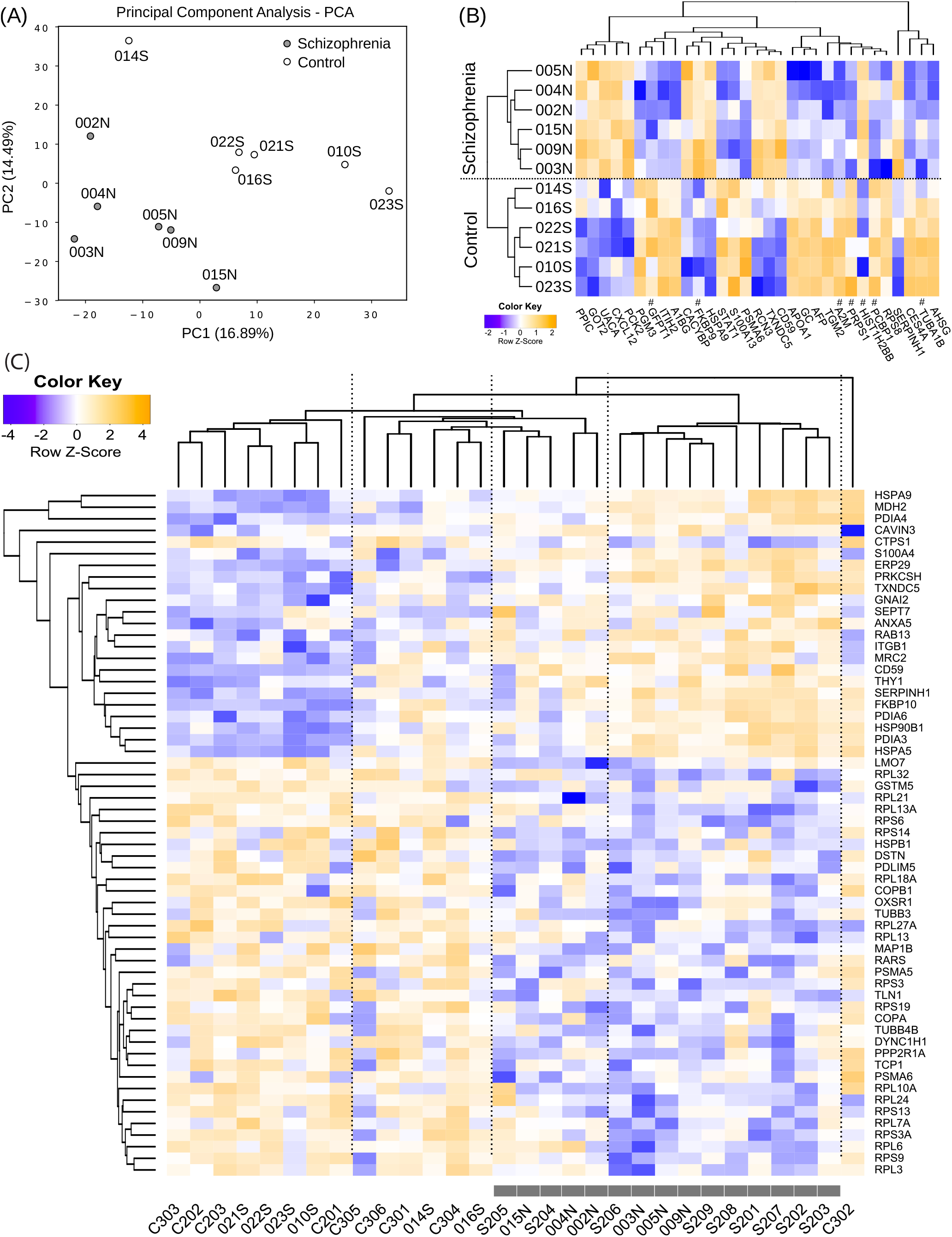
The ONS cell proteome distinguishes adolescent-onset and adult-onset schizophrenia patients from healthy controls. **(A)** Principal component analysis (PCA) of the ONS cell proteome (1638 proteins) shows that adolescent-onset schizophrenia patients (filled circles) clustered according to their disease status and were well separated from adolescent controls (open circles). **(B)** Hierarchical clustering heatmap for differentially expressed proteins in the same patients and controls. #: proteins with other members of the family showing identical peptide profiles. **(C)** The ONS cell proteome from combined analysis of adolescent-onset and adult patients and controls. Hierarchical clustering heatmap for differentially expressed proteins in patients (grey boxes) compared to controls.

### Reduced expression of ribosome, myosin and tubulin proteins in adolescent-onset ONS cells

Two hundred and forty one proteins were differentially expressed (123 down-regulated in patients and 118 up-regulated; p<0.05) (Supplementary Table 1). Forty five proteins remained significantly differentially expressed after multiple testing correction (25 down-regulated and 20 up-regulated; Benjamini-Hochberg adjusted p<0.1; Figure 1B). Eighty five ribosomal proteins were detected in patient and control ONS cells. All detected ribosomal proteins were downregulated in patient ONS cells relative to controls cells, twenty of them significantly (23.5%, p<0.05; Supplementary Table 1). Six out of 14 detected tubulin proteins (42.8%) and six out of 14 detected myosin proteins (42.8%) were also significantly down-regulated in patient cells compared to control cells (p<0.05; Supplementary Table 1).

### Altered protein synthesis and cell adhesion pathways in adolescent-onset ONS cells

Ingenuity Pathways Analysis (IPA) was used to map functional pathways associated with the 241 differentially expressed proteins (Supplementary Table 1). These proteins were over-represented in pathways that can be grouped into broad categories of cell functions (Table 2): protein synthesis and its regulation (“EIF2 signalling”, “Regulation of eIF4 and p70S6K signalling”, “mTOR signalling”); cell adhesion and cytoskeleton (“Epithelial Adherens Junction Signalling”); and metabolism (“PRPP Biosynthesis I”).

**Table 2.** Differentially expressed protein in adolescent-onset schizophrenia. Pathway analysis of differentially expressed proteins in adolescent-onset schizophrenia patients relative to healthy controls (n=12, p<0.05, not FDR corrected) and in combined analysis of adolescent-onset and adult patients and controls (n=30, p<0.05, not corrected for multiple testing).

| **Ingenuity Canonical Pathways** | **-log (p-value)** | **Down-regulated proteins** | **Up-regulated proteins** |
| --- | --- | --- | --- |
| **Adolescent-onset** |  |  |  |
| EIF2 Signaling | 18.7 | RPL24, RPS19, RPS3A, RPS27, RPS8, RPL23, RPL7A, RPS11, RPL28, RPL7, RPL10A, RPS6, EIF3F, RPS16, RPS26, RPS9, RPL21, RPS2, RPL18 | HSPA5, WARS, EIF5 |
| Regulation of eIF4 and p70S6K Signaling | 7.88 | RPS6, RPS19, EIF3F, RPS16, RPS3A, RPS26, RPS27, RPS9, RPS8, RPS2, RPS11 |  |
| mTOR Signaling | 7.83 | RPS6, RPS19, EIF3F, RPS16, PLD3, RPS3A, RPS26, RPS27, RPS9, RPS8, RPS2, RPS11 |  |
| Epithelial Adherens Junction Signaling | 6.11 | MYL9, TUBA1B, TUBB3, MYH9, MYL6, LMO7, TUBB4B, ZYX, TUBB |  |
| PRPP Biosynthesis I | 5.93 | PRPS1L1, PRPS2, PRPS1 |  |
| **Adult-onset** |  |  |  |
| EIF2 Signaling | 32.5 | RPL7A, PTBP1, UBA52, RPL13A, RPS26, RPL32, RPL23, RPL6, RPS19, RPS13, RPL9, RPL3, RPS8, RPL4, RPS3A, RPL14, RPL18A, RPL27A, RPS6, RPL21, RPL13, RPS14, RPS3, RPL7, RPS9, RPL24, RPL34, RPS27A, RPS4X, EIF3F, RPL10A | HSPA5, RRAS |
| Protein Ubiquitination Pathway | 11.9 | UBA52, CRYAB, UCHL1, PSMA6, HSPB1, UBB, RPS27A, HSP90AB1, UBC, PSMA5 | UBE2L3, HSP90B1, USP5, UBE2D3, UBE2D2, HSPA9, PSMB4, HSPA5 |
| Regulation of eIF4 and p70S6K Signaling | 11.9 | RPS6, RPS26, RPS14, RPS3, RPS19, RPS13, RPS9, PPP2R1A, RPS27A, RPS8, RPS4X, RPS3A, EIF3F | ITGB1, RRAS |
| mTOR Signaling | 9.34 | RPS6, RPS26, RPS14, RPS3, RPS19, RPS13, RPS9, PPP2R1A, RPS27A, RPS8, RPS4X, RPS3A, EIF3F | RRAS |
| Epithelial Adherens Junction Signaling | 7.79 | ZYX, ARPC2, MYL6, TUBB3, TUBB, LMO7, MYL9, ARPC5L, RRAS, TUBB4B, MYH9 |  |
| Integrin signaling | 7.04 | ZYX, ARPC2, CAV1, TLN1, ITGB1, ARF4, MYL9, PARVA, ARPC5L, PFN2, CAPNS1 | RRAS |
| Unfolded protein response | 5.44 | VCP | CALR, HSPA9, PDIA6, HSP90B1, HSPA5 |
| RhoGDI signaling | 5.14 | ARPC2, MYL6, ARHGAP1, MYL9, ARPC5L | GNAI2, ITGB1, GNB1, CD44 |

### Similar protein expression in adolescent- and adult-onset ONS cells

Analysis was confined to the 900 proteins that were detected in both adolescent and adult cohorts. The differentially expressed proteins were similar and overlapping in the adolescent- and adult-onset cohorts (148 and 116 proteins in adolescent and adult cohorts respectively, Supplementary Table 1). Across all 900 proteins there was a significant positive correlation in differential protein expression between the two cohorts (Pearson’s correlation coefficient R=0.2, t=6, df=898, p<1e-08). Differential protein expression comparing the adolescent and adult patient cell cohorts revealed only 20 proteins different in the two patient cell cohorts and 45 proteins different in the control cell cohorts, none of which were over-represented in any pathway using IPA analysis. Collectively these observations indicate that the protein expression in the two cohorts was very similar. Therefore the two datasets were combined to increase statistical power with a study size of 30 (15 patients and 15 controls).

### Reduced ribosomal protein expression in adolescent- and adult-onset ONS cells

This combined analysis identified 164 proteins differentially expressed in patient-derived ONS cells (p<0.05; Supplementary Table 1), with 62 proteins significantly different (40.9%; 37 down-regulated and 25 up-regulated) after multiple testing correction (Benjamini-Hochberg adjusted p<0.1). A cluster analysis of differential protein expression classified the patients and controls separately (Figure 1C). Importantly, adolescent-onset and adult-onset patients were clustered together but intermingled. Ribosomal proteins were the largest group of dysregulated proteins in the combined data. All ribosomal proteins were down-regulated in adolescent- and adult-onset patient ONS cells (Figure 2A). Using the combined data we identified 13 ribosomal proteins reaching statistical significance (Figure 2A). A cluster analysis of differential ribosomal protein expression classified the patients and controls into separate groups (Figure 2B).

**Figure 2.**
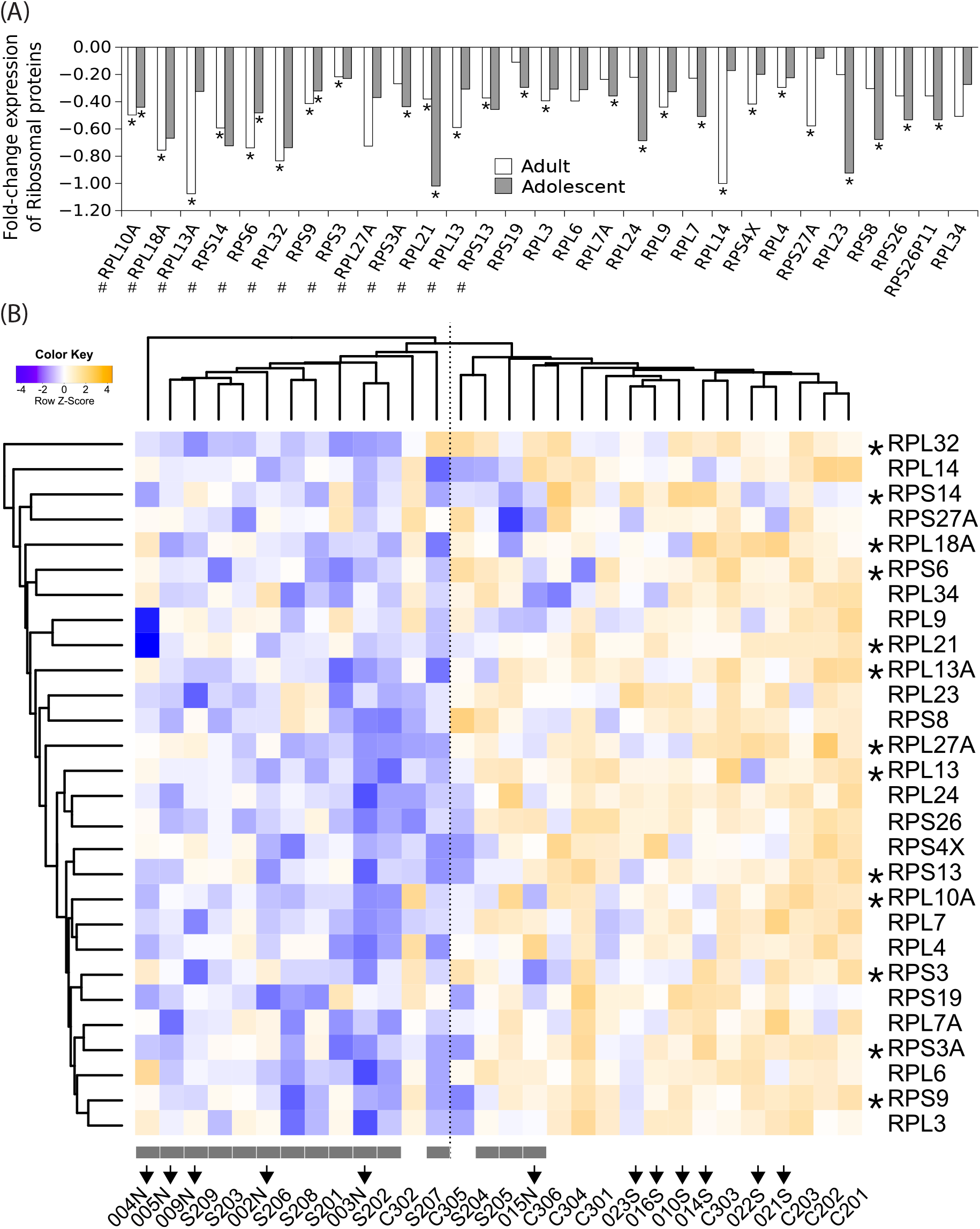
Expression of ribosomal proteins is reduced in schizophrenia ONS cells. **(A)** Fold-change in expression of the ribosomal proteins in two independent cohorts of schizophrenia patients (adolescent-onset and adult-onset) compared to healthy controls. (*) Significant fold-change expression in each cohort (t-test, p<0.05). (#) Significant fold-change expression in combined cohort analysis (t-test, p<0.05). **(B)** Hierarchical clustering heatmap for differentially expressed ribosome proteins in 15 patients compared to 15 controls. Schizophrenia patients from independent cohorts show similar patterns of ribosomal protein expression in ONS cells. Grey boxes indicate schizophrenia patients. Arrows indicate adolescent-onset cohort of patients and controls. *: Statistically different expression between schizophrenia and controls (t-test student p<0.05).

### Protein synthesis and cell adhesion are dysregulated in adolescent-and adult-onset ONS cells

Ingenuity Pathways Analysis of the combined data (Table 2) identified the same altered protein synthesis signalling pathways that were found in the adolescent-onset group: “EIF2 signalling”, “Regulation of eIF4 and p70S6K signalling”, “mTOR signalling”. Disrupted cell adhesion signalling was also replicated (“Epithelial Adherens Junction Signalling” in the adolescent cohort; “Epithelial Adherens Junction Signalling”, “Integrin signalling”, “RhoGDI signalling”) in the combined data which also identified disrupted protein metabolism pathways (“Protein ubiquitination pathway”, “Unfolded protein response”).

We explored the potential coordination between the differentially expressed proteins by generating a protein-protein interaction (PPI) network based on the 164 differentially expressed proteins in the combined dataset. We used these proteins as ‘seeds’ and added their direct (“first-degree”) protein interacting partners present in the ONS cells proteome. This PPI network (ONS-SZ PPI) is comprised of 392 proteins (“nodes” in the network) and 1554 interactions (interconnections between nodes in a network, “edges”) in which 146 (89%) of the 164 differentially expressed proteins are present (Supplementary Figure 1). The differentially expressed proteins in the patient cells were highly interconnected, with the statistically identified proteins (seeds) distributed throughout the network (Supplementary Figure 1). We modelled the interconnectedness of the network by simulation. By all measures, the ONS-SZ PPI network was significantly different from the random simulation with all measures indicating a significantly more interconnected network (Figure 3A-C). In the ONS-SZ PPI network the average degree (number of other nodes a given node is connected to) was significantly greater than random (D_ONS_=5.3 vs D_random_=3.1; p=2.2e-16, Kolmogorov-Smirnov test); the average path length (average of number of nodes between all pairs of nodes in the network) was significantly less than random (L_ONS_=3.5 vs L_random_=4.5; p=2.2e-16); and the average clustering coefficient (a higher-order measurement to quantify how well neighbours are interconnected) was significantly greater than random (C_ONS_=0.017 vs C_random_ =0.0005, p=2.2e-16). The high degree, short path length and high clustering coefficient of the ONS-SZ PPI network strongly suggests that differentially expressed proteins and their partners are physically connected and working in concert to regulate functional pathways and cell functions. The ONS-SZ PPI network is dominated by the ribosomal proteins and their interconnections (Figure 3D). The other dominant modules, linked by ribosomal proteins, represent regulation of protein synthesis (“mTOR signalling”, “EIF2 signalling”) and cell adhesion (“Adherens junction signalling”), thus replicating the significant pathways identified in the adolescent-onset cohort (Table 2).

**Figure 3.**
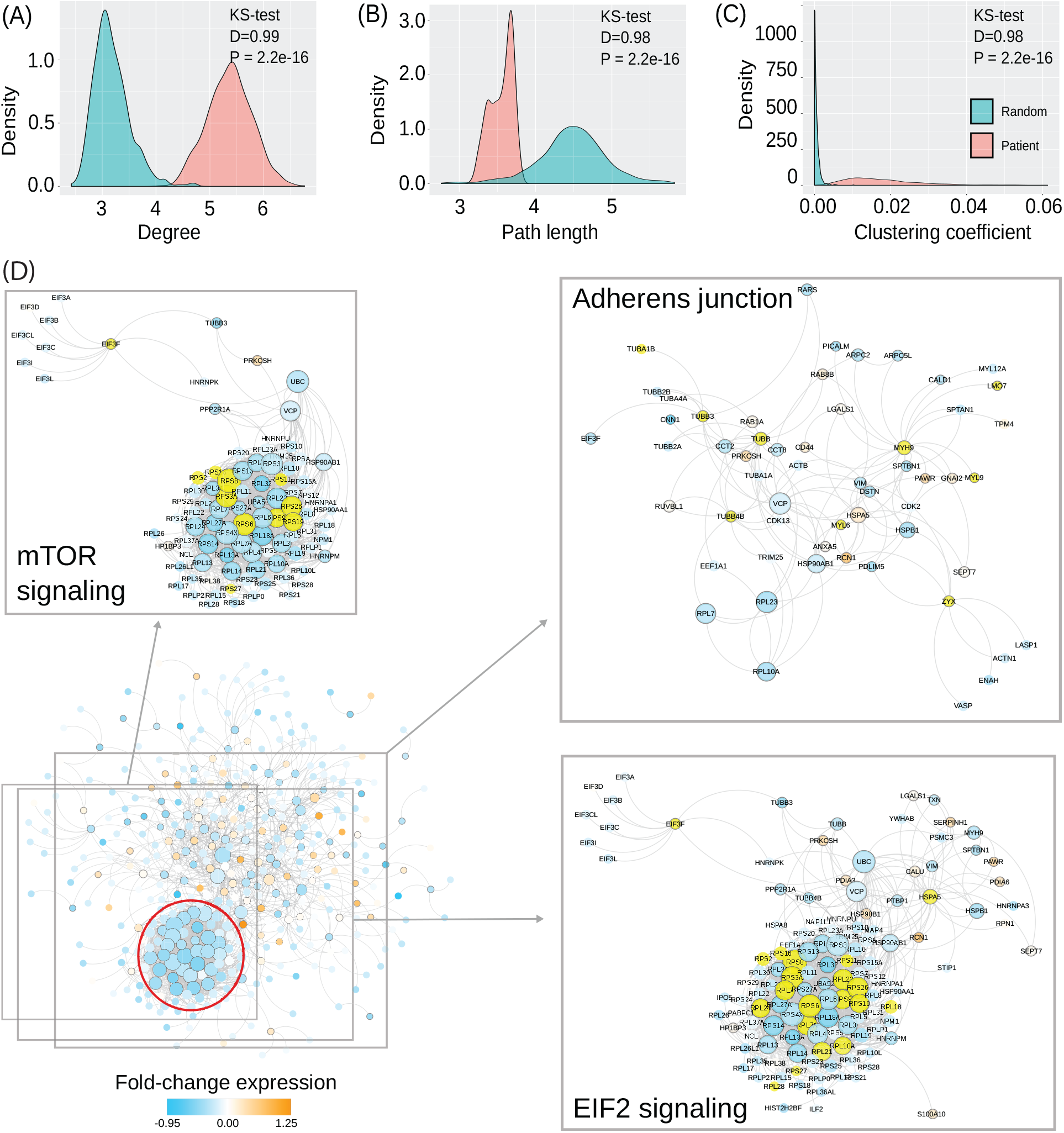
ONS-SZ PPI network proteins are significantly more interconnected than in randomly generated PPI networks. The degree of interconnectivity **(A)** is higher, the path length between nodes **(B)** is shorter, and the clustering coefficient of the network is higher **(C)** in ONS-SZ PPI network (pink) than randomly generated networks (turquoise). **(D)** High levels of interconnectivity reveal protein interaction modules in ONS-SZ PPI network. Three enriched functional groups are showed in more details with proteins identified by IPA pathway analysis highlighted in yellow. The red circle surrounds the subnetwork containing the ribosomal proteins.

### Relevance of patient ONS cell proteome to schizophrenia and neurodevelopment

We tested how well the ONS-SZ PPI network (based on protein expression in ONS cells) overlaps with the AXAS-SZ PPI network (based on genetic variation and association in the population): these PPI networks overlapped significantly more than expected by chance alone (Z-score=7.29; P=2e-13). The AXAS-SZ PPI network was built from published studies of schizophrenia-associated protein-coding genes ^24^ and their known protein-protein interactions ^21^. We then tested how well the ONS-SZ PPI network overlaps with other schizophrenia-associated proteome and transcriptome datasets (Supplementary Table 2). There was significant overlap with the ONS-SZ PPI network in: 1) protein expression in schizophrenia patient-derived neuronal progenitors generated from induced pluripotent stem cells ^25^ (Z-score=16.61; P=3e-62), 2) RNA expression in patient-derived ONS cells (Z-score=3.24; P=0.0004; ^5^), 3) RNA expression in patient-derived lymphoblastoid cell lines (Z-score=2.75; P=0.003; ^26^), and 4) variants in expressed genes associated with schizophrenia (Z-score=1.96; P=0.03; Opentargets database ^27^). There was no overlap with RNA expression in schizophrenia patient-derived fibroblasts (Z-score=−0.27; P=0.6, ^5^).

We next tested how well the ONS-SZ PPI network overlaps with genes involved in neural stem cell differentiation in mouse brain ^28^. There was significant over-representation in the ONS-SZ PPI of genes that were upregulated as neural stem cells differentiated into neuroblasts (Z-score=2.22; P=0.01) and downregulated as neuroblasts differentiated into neurons (Z-score=8.58; P=5e-18; Supplementary Table 2).

We then tested how well the ONS-SZ PPI network overlaps with genes expressed in axon growth cones in the of the corpus callosum in the developing mouse forebrain ^29^. The growth cone associated RNAs and proteins were significantly over-represented in the ONS-SZ PPI network (Z-score=26.85; P=5e-159) including proteins involved in cell adhesion and in protein synthesis (Supplementary Table 2).

Finally, we tested how well the ONS-SZ PPI network overlaps with a PPI network based proteins associated with the focal adhesion complex ^30^. This complex links the integrin receptors (which detect the extracellular matrix) and the actin cytoskeleton (though which cells exert force) to regulate cell adhesion and motility. Adult schizophrenia-patient derived ONS cells are less adhesive than control cells ^8, 10^ and are dysregulated in transcriptome, proteome and functions related to cell motility and axonal outgrowth ^5, 8, 10, 11, 31^. Focal adhesion PPI network proteins were significantly over-represented in the ONS-SZ PPI network (Z-score = 9; P = 1e-19; Supplementary Table 2).

### mTOR signalling is downregulated in schizophrenia ONS cells

The reduced expression of ribosomal proteins suggested that mTOR signalling is repressed in the patient ONS cells. Additionally, mTOR signalling was identified as a dysregulated pathway and represented as a highly interconnected node in the ONS-SZ PPI network. Therefore we investigated whether the differentially expressed proteins identified in schizophrenia patient-derived ONS cells showed mechanistic evidence of repressed mTOR activity by searching for the canonical 5’-terminal oligopyrimidine-rich (TOP) motif (CCTCTTTTCCG) in the 5’-UTR regions of down-regulated proteins in our schizophrenia patient-derived ONS cells. mTOR stimulates translation of proteins with TOP motifs in their mRNA, including protein translational machinery (several initiation and elongation factors, and many ribosomal proteins) ^32^. Transcripts of four down-regulated proteins in patient ONS cells have highly conserved TOP motifs (RPS6, VIM, LDHB and PPP2R1A) and we found TOP motifs significantly enriched in down-regulated proteins (Fisher’s exact test P=3.7e-06) but not in up-regulated proteins. Combined with the coordinated reduction in ribosomal protein expression the reduction in expression of TOP motif proteins strongly suggests that mTOR activity is repressed in the patient cells.

## Discussion

This discovery-based proteomics study has revealed protein expression that is dysregulated in patient cells from adolescent-onset schizophrenia compared to age- and gender-matched healthy controls. Ribosomal proteins were particularly affected: 14% (241/1638) of the identified proteins were differentially expressed in the patient cells, with the 85 ribosomal proteins downregulated. The dysregulated proteins were significantly over-represented in cell signalling pathways concerned with protein synthesis and its regulation, cell adhesion, and cytoskeleton. These findings essentially replicated our previous discoveries in ONS cells from adult-onset schizophrenia ^11^. The two studies were independent, separated by patient and control populations, location and time, but linked by the same cell model grown and maintained under rigorous handling and culture protocols. In spite of these potential confounds the patients from these groups were classified together in cluster analyses of global and ribosomal protein expression differences, distinct from the controls. The present study of adolescent-onset schizophrenia ONS cells thus replicates and validates the protein expression first reported in ONS cells from adult-onset schizophrenia ^11^. When the two data sets were combined to increase statistical power (n=30 in combined analysis), 164 proteins were significantly differentially expressed. Ribosomal proteins were universally down-regulated, with 13 reaching statistical significance, and pathway analysis identified the same significantly dysregulated functions: protein synthesis and its regulation, cell adhesion and cytoskeleton. Protein ubiquitination and the unfolded protein response were also significantly dysregulated. These results support hypotheses of coordinated dysregulation of cell functions shared by individual patients, despite different populations, age, different medication histories, and genetic risk profiles ^21, 33–35^. A protein-protein interaction (ONS-SZ PPI) network was constructed from the differentially expressed proteins in the combined data. This highly interconnected ONS-SZ PPI network demonstrates a remarkable degree of coordination in the dysregulation of protein expression in the patient ONS cells (89% of the differentially expressed proteins were connected by the network). It reveals that the majority of the differentially expressed proteins formed clusters, network modules, around “mTOR signalling”, “EIF signalling” and “Adherens junction”. This ONS-SZ PPI network was significantly more interconnected than random as evidenced by computational analysis of simulated networks based on randomly chosen proteins. In summary, this study demonstrates that ONS cells from patients with adolescent-onset schizophrenia have significantly reduced expression of ribosomal proteins and related proteins that overlap with protein expression in adult-onset schizophrenia. Bioinformatic analysis identified similarly affected cellular regulatory pathways in adolescent-onset and adult-onset patient cells. Network analysis of the combined dataset indicates significant co-regulation of sub-networks of proteins in the patient cells that are consistent with reduced protein synthesis, demonstrated in the adult-onset cohort ^11^.

We previously generated the AXAS-SZ-PPI network from a curated list of schizophrenia risk genes, a sub-network of the AXAS-PPI network of genes involved in several neurodevelopmental diseases ^21^. Individuals are unlikely to carry all the genetic risks identified in the general population but probably have a subset of mutations that converge on regulatory networks that control cell functions similarly across individuals. This working hypothesis was confirmed by the significant overlap of the ONS-SZ-PPI network with the AXAS-SZ-PPI network, indicating that regulatory networks of protein expression in affected individuals (ONS-SZ PPI network) converge on regulatory networks identified in population genetic risk (AXAS-SZ-PPI network). Further evidence for this hypothesis was demonstrated in the epigenome and transcriptome of patient-derived ONS cells: schizophrenia-associated DNA methylated gene loci and gene expression were also significantly associated with the AXAS-SZ-PPI network ^9^. These observations suggest that patient-derived ONS cells carry genetic risks identified in large population studies and will be a useful model to identify the relations between genetic risks and functional consequences in individuals with schizophrenia.

What conclusions can be drawn about schizophrenia, a brain disease, from ONS cells, a neural stem cell derived from olfactory mucosa? Schizophrenia is widely accepted as a neurodevelopmental disease arising from early events that affect brain development ^2, 3^. Genetic and environmental risk factors are predicted to affect multiple cellular processes that build brain structure ^36^. How do the many schizophrenia-associated ONS cell phenotypes, including proteomes, fit into concepts of brain development and function? These ONS cell phenotypes are all associated with brain development: reduced protein synthesis ^11^, reduced adhesion, altered focal adhesion dynamics and dysregulated motility ^8^, lack of motility response to extracellular matrix proteins, including reelin ^10, 31^, smaller cell size and reduced expression of cytoskeletal proteins ^10^ and faster cell proliferation ^7^. Any of these dysfunctions could alter the trajectory of brain development and many are functionally inter-connected. For example, cell adhesion regulates protein synthesis, cell size and morphology, cell proliferation and motility ^37^. In the growth cones of developing mouse cerebral cortex neurons, ribosomal and cytoskeletal RNAs and proteins are enriched ^29^ and we show here they overlap significantly with ONS-SZ PPI network modules involved in protein synthesis, including mTOR and RNAs that contain mTOR motifs, ribosomal proteins, and cytoskeletal components ^29^. Axon growth and neuron migration depend on cell adhesion to the extracellular matrix via focal adhesions, multi-protein complexes at the cell surface that link the extracellular substrate to the actin-cytoskeleton to control cell adhesion, cell morphology and cell motility ^30^. Focal adhesion complexes are enriched in ribosomal proteins, proteins involved in protein synthesis and cytoskeletal proteins ^30^ and we show here the significant overlap of the focal adhesion complex proteins with the ONS-SZ PPI network. Therefore the dysregulation of protein expression, focal adhesions and protein synthesis in patient ONS cells can be directly related to molecular mechanisms in growth cones and the growth of axons during brain development.

Protein synthesis is very dynamic during neurogenesis: it is low in neural stem cells, rises in neural precursors and drops again after differentiation into neurons ^28^. We show here significant association of the ONS-SZ PPI network with the transcriptomes of neural stem cells as they pass along this neural differentiation route. During brain development, protein synthesis in cortical neurons is reduced via suppression of mTOR signaling ^38^. Patient ONS cells have reduced protein synthesis 11 and downregulated expression of protein synthesis machinery (e.g. ribosomal proteins, RPS6, EIF3F) indicating reduced activation of mTOR ^39^. Activation of mTOR phosphorylates the ribosomal protein RPS6 (whose expression is less in patient ONS cells) and stimulates translation of certain growth-related mRNAs, including several initiation and elongation factors, and some ribosomal proteins which are defined by a 5′-terminal oligopyrimidine (TOP) motif recognised by the eukaryotic initiation factor 4F (eIF4F) ^32^. In patient ONS cells, proteins with TOP motif mRNAs were significantly down-regulated, suggesting suppression of mTOR activity. Four proteins with the canonical TOP motif were significantly reduced in expression in patient-derived ONS cells compared to controls (RPS6, VIM, LDHB and PPP2R1A).

Dysregulated mTOR signalling is also indicated in other studies. In post mortem brain, ribosomal RNA was increased in the dorsal raphe nucleus in a subset of schizophrenia patients ^40^. In schizophrenia patient leucocytes ribosomal RNA was increased ^41^ however in lymphoblastoid cell lines ribosomal RNA was downregulated when exposed to dopamine ^42^. In contrast tour result using ONS cells, in iPS-derived neural precursors from schizophrenia patients ribosomal proteins were increased ^25^. Taken together these reports suggest that mTOR function and hence protein synthesis may be less stable in schizophrenia, varying with cell type and age. This instability could affect brain development and adult brain function because mTOR regulates focal adhesion and actin cytoskeleton formation to affect cell adhesion, axon outgrowth and synaptic stability ^43–46^. Our findings of the same dysregulated protein synthesis pathways in adolescent- and adult-onset patient cells places our findings at the heart of brain development by providing a site where schizophrenia-associated dysregulation could affect the growth of axons and dendrites and alter the trajectory of brain development.

The adolescent-onset proteome replicates our published adult-onset study ^11^. Acknowledged variability between such replication studies include: locations of laboratories and teams, and differences in protocols ^47^. Despite potential technical differences between the studies, the adolescent-onset cohort confirmed the protein expression differences in the adult-onset cohort, giving confidence that conclusions drawn about the schizophrenia-associated differences are reproducible and validly generalised. Such small case-control studies are open to selection bias in both patient and control samples. We confirmed that there were no control-control, nor patient-patient differences in protein expression between the studies. Thus the patient and control samples could be combined to increase the statistical power and confidence in the findings (combined study size of 30). Some potential sampling confounders can be hard to control (age-of-onset, smoking, medications, period-of-medication). The similarity in protein expression between adolescent and adult cohorts indicates that differences in these potential confounds were unlikely to affect our conclusions. Finally, a caveat: the number of schizophrenia-associated differences in protein expression is probably underestimated because only the most abundant proteins are likely to be detected and identified in global proteomics analysis ^48, 49^. These include the ribosomal and cytoskeletal proteins that figure so prominently in our analysis.

In summary, the proteome of adolescent-onset patients with schizophrenia provides evidence for repression of protein synthesis via mTOR, indicated by the universal downregulation of ribosomal proteins and of proteins with TOP motif mRNA. We also find a remarkable similarity of the proteome of ONS cells from adolescent-onset and adult-onset schizophrenia patients, notably in proteins associated with protein synthesis, cell adhesion and cell migration. A PPI interaction network built from the differentially expressed proteins was highly co-ordinated and tightly interconnected. This ONS-SZ-PPI overlapped significantly with PPIs based on the focal adhesion complex, neural precursors and axon growth cones in the developing mouse cortex. Although the present data suggest similarity in the aetiology of schizophrenia in adolescent- and adult-onset patients, it will require larger samples and deeper protein coverage to confirm this. Future studies should also focus on the mechanisms of mTOR dysregulation in schizophrenia.

## Acknowledgements

Thanks to the participants for their involvement and their olfactory tissues. Thanks to Dr Pablo Devesa for collecting the biopsies, Prof Olaf Ansorge for providing the human tissue culture facilities, Dr Clare Bosay for undertaking the psychometry and Miss Kristina Roth for assessing the cellular behaviours. Access to and use of mass spectrometry instrumentation and computing facilities at the Conway Institute is gratefully acknowledged. This work was supported by the Irish Health Research Board through a Health Research Board Clinician Scientist Award (to DRC) and the 2013 BMA Margaret Temple Award (to AJ).

